# A network optimisation condition uncovers the role of functional groups in the feasibility and dynamical stability of microbial model ecosystems

**DOI:** 10.1101/2024.01.09.574826

**Authors:** Léo Buchenel, Sebastian Bonhoeffer, Alberto Pascual-García

## Abstract

Investigating mechanisms favouring stable coexistence of complex microbial communities is central to understand their formation and maintenance in the natural environment and, eventually, controlling and designing synthetic communities. In this work, we studied microbial model ecosystems in which consumption and secretion of resources was explicit, determining ecological interactions (e.g. competition for resources, facilitation, mutualism). We found a sufficient condition for the dynamical stability of microbial model ecosystems. This condition allowed us to derive an objective function whose optimization selected specific configurations of consumption and secretion of resources. Optimized configurations stood out as a compromise between having a large parameter space in which species coexistence was feasible (dominated by systems in which microbes do not secrete any resources), and having a high rate of return to equilibrium after a perturbation (dominated by systems in which every microbe secrete all resources). We explained the behaviour observed for optimized configurations by noting that they host sets of species with differentiated niches -as defined by their consumption and secretion strategies-, termed functional groups. We speculated that, since increasing the number of functional groups increased the number of niches, competition between species should be reduced, favouring coexistence. Therefore, our results suggest that the formation of functional groups have an important role in microbial coexistence, which has ramifications for the design of complex microbial communities.

---

The importance of microbial communities (MCs) is increasingly recognized. For instance, their control may have enormous benefits for human health [1], bioremediation [2, 3], land use [4], biofuel production [5], or lignin degradation [6]. However, our understanding of the complexity of the ecological and evolutionary mechanisms governing assembly and functioning of MCs is still in its infancy. Many important functions in which MCs are involved occur in time-scales much longer than bacterial generation times and, hence, it is of great interest to understand the determinants of MCs’ stability.

In macroscopic organisms, a large body of theoretical research has been developed around Lotka-Volterra (LV) models, aimed at understanding the relative influence of ecological interactions on the stability of ecosystems. With these models, new concepts have emerged explicitly linking ecological interactions with the determinants of coexistence, such as storage effects [7], the connection between the effective competition in the system and its global stability [8], or the influence of interaction patterns such as the nestedness [9]. Also novel angles to study the stability of ecosystems beyond dynamical stability were developed, such as structural stability [10, 11] or permanence [12].

The dynamics of microbial communities, however, is strongly influenced by the available nutrients in the environment which may fluctuate in very short time-scales. As a consequence and, although some agreement with experimental data under laboratory conditions was achieved with LV models [13], it is expected that explicit modelling of the resources’ dynamics is needed to achieve predictions under natural conditions [14]. Following the seminal theoretical work of MacArthur [15] and Tilman [16], the surge of experiments involving microbes has prompted renewed interest in consumer-resource models.

More recent theoretical work showed that microbial communities can shape their own environment to achieve coexistence, even if no syntrophy was present in the system [17] or if there were more species than resources [18], under the assumption that resources equilibrate in time-scales shorter than the population dynamics. Moreover, analysis of cohesive communities led to the identification of optimal metabolic classes an how their combination explain stable strategies [19]. Extending these results to models in which the dynamics of resources is not at a steady-state, and in which feedbacks between microbes and the environment exist through the release of by-products, represents a challenge [20]. The release of by-products of some species that can be consumed by others is a form of mutualism, termed syntrophy, widespread in the microbial world [21]. LV models predict an important role for mutualistic interactions in biodiversity maintainence [22], with a complex interplay with competitive interactions [11]. However, the relative role of competitive and mutualistic interactions in consumer-resource models is less well understood, and specially the role of the topology in networks describing the consumption and secretion of metabolites.

In this work, we considered a consumers-resources model with syntrophy, and performed an exhaustive exploration of the feasibility and dynamical stability of this system. We found that syntrophy hinders feasibility but enhances dynamical stability. In addition, we show that it is possible to find a sufficient condition for dynamical stability, which has so far remained elusive [23]. The condition allowed us to propose an optimization algorithm to design interaction matrices increasing the likelihood of being dynamically stable, i.e. increasing the volume of feasible and dynamically stable fixed points. Finally, we show that the mechanism explaining an increase in the feasibility of optimized systems in turn relies on an increase -with respect to their random counterparts-in the number of functional groups.

We considered that the dynamics of population densities *S*_*i*_ of *N*_S_ species and of resources *N*_R_ consumed and secreted by microbes, *R*_*μ*_, are governed by the following set of differential equations [23]:

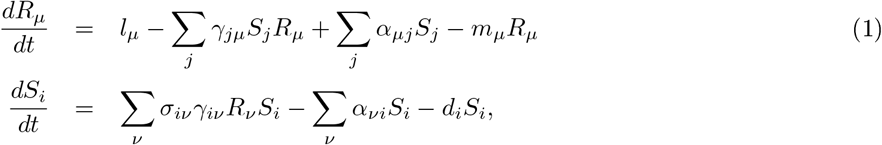

where the non-negative matrices *γ*_*jμ*_ (*α*_*μj*_) describe the uptake (secretion) rate of resource *μ* by species *j*, and we considered continuous growth under a constant supply of resources *l*_*μ*_, and dilution rates *m*_*μ*_ and *d*_*i*_ for resources and species, respectively. We further considered that only a fraction *σ*_*μ*_ of the resources consumed are converted into biomass. Note that the secretion of resources only depends on the densities of species and not on the resources they consume. In the original formulation of the model this was interpreted as an appropiate choice for by-products of central metabolism, and/or biomass released in the environment upon cell death [23]. The model also imposes conservation of biomass by subtracting the biomass released by the organism from their biomass, and we impose the following conservation of biomass condition to hold

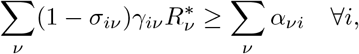

which ensures that the biomass secreted is smaller or equal to the fraction not converted into biomass at steady state *R*^*\**^.

The characteristic equation of the Jacobian of the system has no explicit solution, as detailed in Supplementary Materials. With the help of the Gerschgoring circle theorem [24] it is, however, possible to find an upper bound (hereafter critical radius *r*_c_) for the modulus of every eigenvalue of the Jacobian, i.e. |*λ*| ≤ *r*_c_ ∀*λ*. This allowed us to show that if 0 is not an eigenvalue of the Jacobian at a given equilibrium, and the following condition holds for every resource:

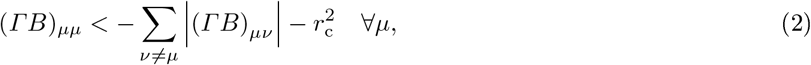

then the system is dynamically stable (see SM for proof and explicit definition of *r*_c_). Note that the product of matrices Γ = *γ*diag(*R*^*\**^) + *α* and B = diag(*S*^*\**^)*σγ* imposes constraints on the relation between the consumption matrix *γ* and the secretion matrix *α*, and that the sufficient condition expresses that the matrix ΓB is diagonally dominant. By working with mean field matrices *γ* = *γ*_0_G and *α* = *α*_0_A (where *γ*_0_ and *α*_0_ are metaparameters drawn from uniform distributions and G and A are binary matrices) and, after some simplifications detailed in Supplementary Materials, we found that it is possible to find an objective function whose minimization increases the likelihood of finding dynamically stable systems:

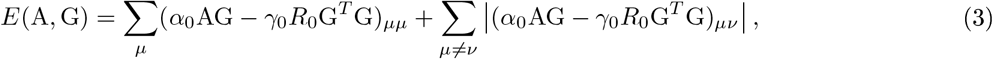

which generalizes the condition found in [23] for systems hosting only specialist consumers. We studied the topologies obtained by minimizing the condition in Eq. 3, by considering a large set of consumption matrices with *N*_R_ = *N*_C_ = 25 with different connectances and mean ecological overlap, under a wide range of values detailed in Supplementary Materials (Suppl. Fig. 1). The ecological overlap measures the consumption (secretion) overlap of a pair of species *i* and *j* by counting the fraction of resources consumed (specified in matrix *X* = G) or secreted (specified in *X* = A) by both species, i.e. ∑_*μ*_ *X*_*iμ*_*X*_*jμ*_*/* min(∑_*μ*_ *X*_*iμ*_, ∑_*μ*_ *X*_*jμ*_). We then defined the mean consumption (secretion) overlap, *η*_G_ (*η*_A_) taking the mean across all pairs.

We searched with a Monte Carlo algorithm matrices *A*_opt_ minimizing the objective function in Eq. 3 for each consumption matrix G (Fig. 1A) and by fixing *α*_0_ = *γ*_0_ = *R*_0_ = 1. Since, for different Monte Carlo runs lead to slightly different *A*_opt_ matrices, we adopted the following approach detailed in Supplementary Materials: For each consumption matrix G, we sampled a set {*A*_opt_} and generated a metamatrix *p*_*μj*_ quantifying the probability of having a non-empty cell in position *μ, j* equal to its observed frequency in {*A*_opt_}. We then generated for each consumption matrix: i) a random matrix A_OM_ (optimized ‘metamatrix’) according to the correspondent metamatrix probabilities; ii) random matrices A_RM_ having the same mean connectance than the correspondent A_OM_ as the only constraint; and iii) fully connected syntrophy matrices (A_FCM_, FCM systems). Note that our approach is conservative since we do not directly work with optimized matrices, but the cost is that we might be losing potentially relevant constraints acting on sets of matrix cells.

**Figure 1:**
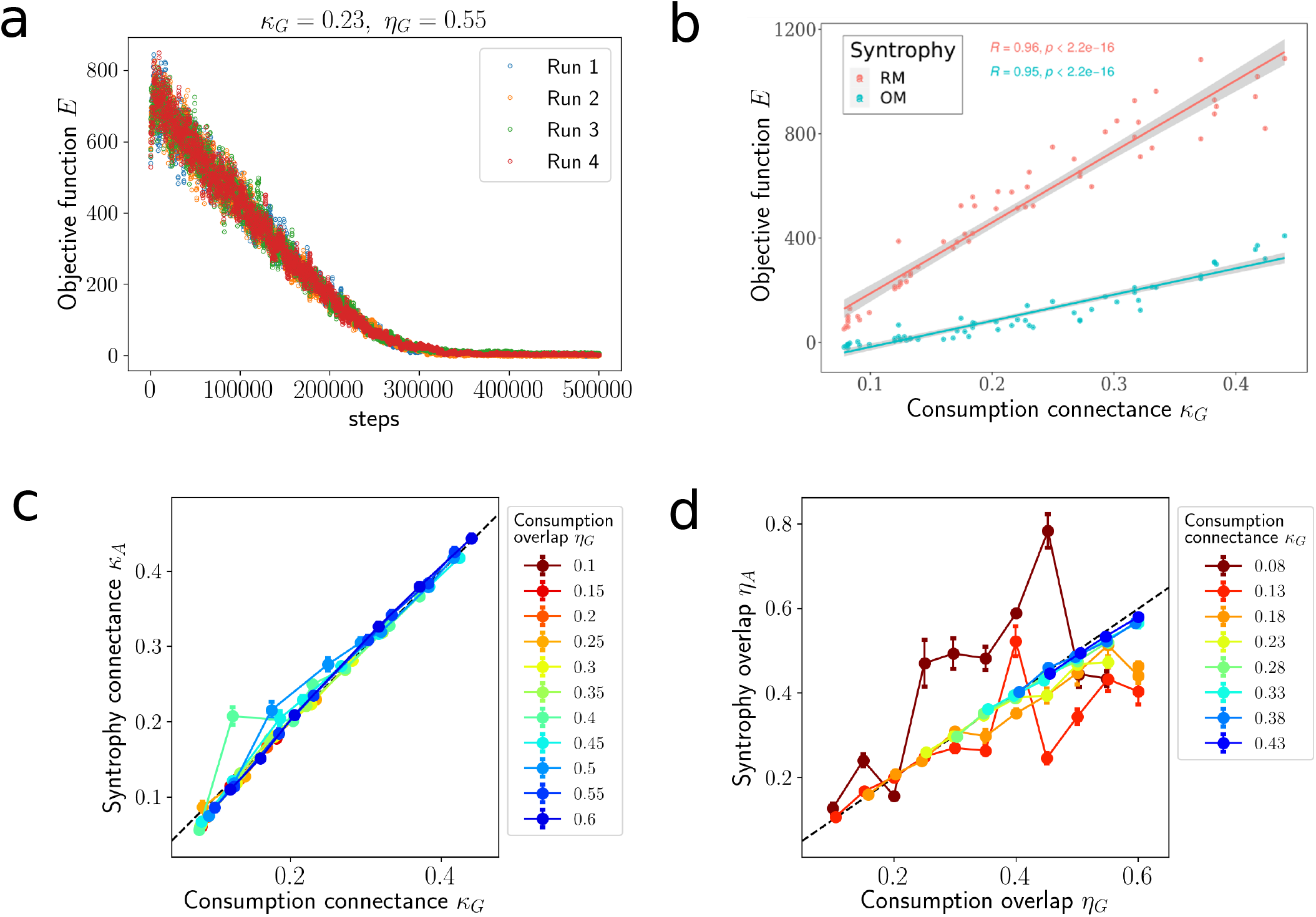
(a) Example of minimization of *E* for a system with consumption connectance *κ*_*G*_ = 0.23 and ecological overlap *η*_G_ = 0.55. Different runs are displayed. The equilibrium is attained within a narrow range of *E* in which different configurations are visited. These configurations are sampled and the probability of observing a syntrophy link estimated. These probabilities are used to draw the set of OMs. (b) Final value of *E* for systems with different consumption topology and a RM syntrophy (red) or OM syntrophy (blue). The higher the connectance the more effective the minimization. (c) Optimized syntrophy matrices have similar connectance and ecological overlap than their correspondent consumption matrices while it is not always the case for the ecological overlap (d), which shows a more erratic behaviour for systems with low consumption connectance, were the minimization is less effective.

The specific value of the minimization depended on the connectance of the consumption matrix G (Fig. 1B and Suppl. Fig. 2) with higher values found for higher connectances. By comparing these values with randomly generated matrices *A*_RM_ with the same connectance, we observed that the minimization is more effective for highly connected matrices (Fig. 1B). Interestingly, the connectance of the optimized syntrophy matrices matched the one of the consumption matrix (Fig. 1C). This might be expected, since both terms in Eq. 3 cancel out for the parameterization chosen. For the ecological overlap, however, the outcome was more unpredictable for low connectance matrices (Fig. 1D), possibly because the searching space is larger and the effect of the minimization less pronounced.

To make systems with different A matrices comparable, we set for all systems the same metaparameters (*γ*_0_, *α*_0_, *l*_0_, 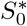 and 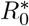, see Suppl. Table 1), and we verified the feasibility of each system by solving the ODEs at the fixed point for *m*_*μ*_ and *d*_*i*_, and asking if these parameters were positive (see Supplementary Materials for details and Suppl. Fig. 3 for an example). For each combination of metaparameters, we generated 200 realizations by randomly drawing parameters from uniform distributions around the metaparameters, and we considered a combination feasible if all realizations were feasible. Subsequently, we asked if the feasible fixed point is dynamically stable by linearizing the system and verifying that all real parts of the eigenvalues of the Jacobian were negative. We considered that a system was linearly stable at each combination of metaparameters if all the realizations were linearly stable.

For all pairs of G and A matrices we found that increasing the syntrophy strength *α*_0_ reduced the volume compatible with feasible and dynamically stable systems (Fig. 2A). Results were qualitatively equivalent if only feasibility was imposed (Suppl. Fig. 4) suggesting that feasibility implied dynamical stability. More specifically, systems with no syntrophy (*α*_0_ = 0) had a larger dynamically stable volume. This was expected since non-zero syntrophy imposes more stringent constraints to solve the ODEs at a fixed point. Consistent with this idea, the volume where dynamical stability was fulfilled was smaller for FCM systems, and it decayed faster with increasing *α*_0_. On the other extreme, OM led to larger volumes than FCM and RM on average (Fig. 2A and Suppl. Fig. 4).

**Figure 2:**
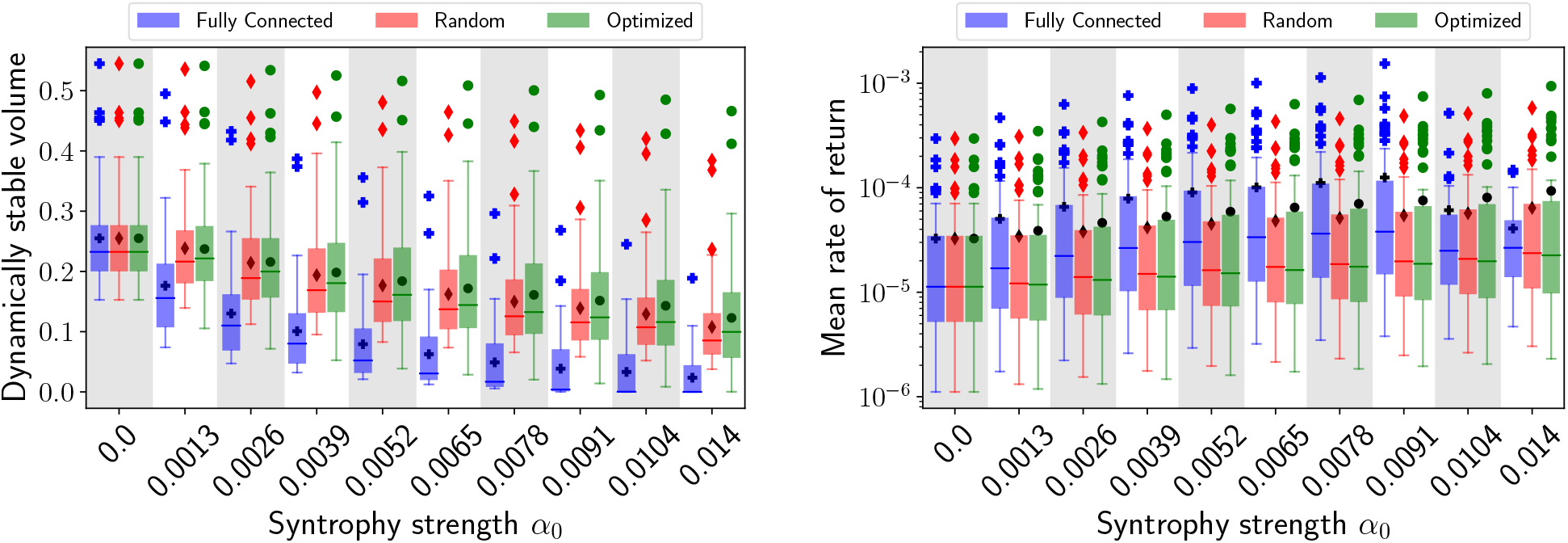
Analysis of dynamical stability. (A) Decay of the dynamically stable volume for different syntrophy strengths and syntrophy matrix topologies. (B) Rate of return to equilibrium after a perturbation in the abundances, versus the syntrophy strength for different topologies of the syntrophy matrix. The limits of each box determine the interquartile region, and the line indicates the median. Means (outliers) are represented with black (coloured) symbols.

Next we analysed, for those systems that were dynamically stable, the rate of return to equilibrium of resources and species abundances after a perturbation, estimated as the absolute value of the dominant eigenvalue of the Jacobian, averaged throughout feasible realizations. We found that syntrophy enhanced the rate of return, and that this increase was more marked for FCM systems (Fig. 2B). Neverheless, when syntrophy was high, since the likelihood of falling into unfeasible parameterizations increased for FCM systems, there was an abrupt drop in the rate of return, i.e. the remaining (feasible) systems had a lower rate of return. On the other hand, the median rate of return for OM and RM systems was comparable, with OM systems having a higher mean throughout the entire range of syntrophy values tested. These findings suggested that OM systems stood out as a compromise between feasibility and recovery after perturbations.

To understand the mechanism explaining this behaviour for OM systems we noted that the minimization of *E* corresponded to configurations in which species had different ecological strategies, namely differentiated patterns of consumed and secreted metabolites, hence exploiting different niches. This point is illustrated in Fig. 3A for a simplified example of two resources and three species. Notably, this differentiation is aligned with the Eltonian definition of guild [25] and, in the context of microbes, of functional groups [26]. To explore whether the existence of functional groups had an influence in our results we identified, for each OM and its correspondent RM system, their functional groups with the algorithm functionink [26]. In brief, functionink is an unsupervised community detection method that allows for the identification of functional groups in multiplex networks. It considers a definition of guilds that departs from a similarity metric quantifying, for each pair of species (metabolites), the number of neighbours shared having the same type of connection (consumption or secretion for species, consumer- or secretory-species for metabolites). Then performs an unsupervised search for a threshold on the similarity determining the optimal partitioning of elements in functional groups (see Suppl. Fig. 5).

**Figure 3:**
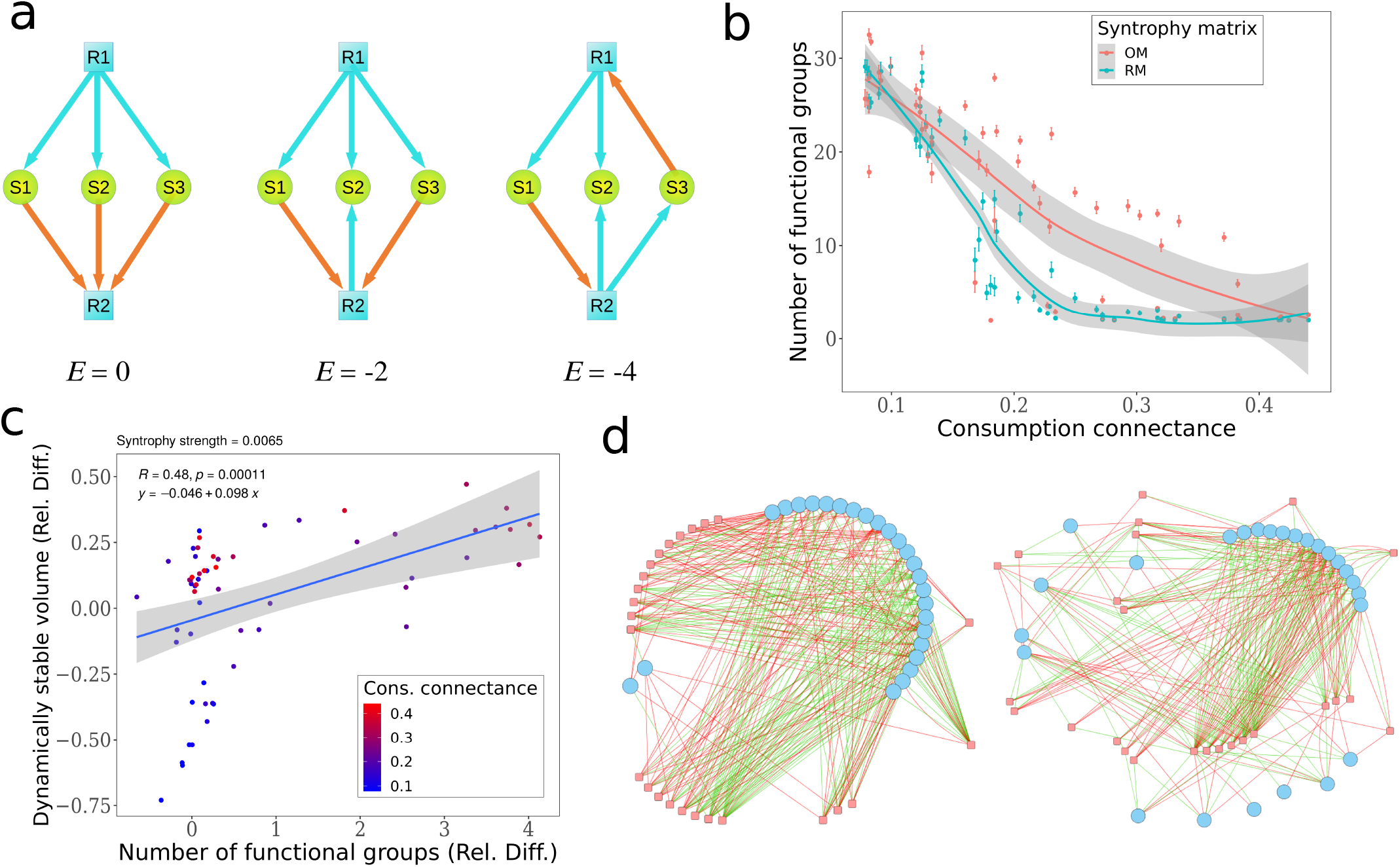
(a) Illustration of a simplified system of three species (circles) and two resources (squares). Depending on the consumption and secretion strategy of each species, the value of *E* varies, with the minimum attained when the three species have different strategies, determining three functional groups. (b) Number of functional groups identified by functionink for systems with different consumption matrices and RM syntrophy (blue) or OM syntrophy (red). The increase is more nuanced for intermediate densities. (c) Relative diference in the volume where dynamical stability was verified between OM and RM systems with the same consumption connectance, versus the relative diference in the number of functional groups. Positive values indicate higher values for OM systems. An intermediate syntrophy strength (*α*_0_ = 0.065) was selected. The positive trend is significant, although it does not have a positive effect for systems with low connectance. (d) Network of a system with consumption connectance *κ*_*G*_ = 0.23 and ecological overlap *η*_*G*_ = 0.55 for RM syntrophy (left) and OM syntrophy (right) with the same connectance. Species are displayed as circles and resources as squares, with red lines indicating consumption and green lines secretion. Nodes belonging to the same functional group are close in space. Despite of the density of the network, a notable increase in the number of functional groups is apparent for the OM system.

We found that the number of functional groups increased for OM systems with respect to equivalent RM-systems for intermediate connectances of the consumption matrix (see Fig. 3B), where the minimization was more nuanced (Fig. 1B). Moreover, by computing the relative increase in functional groups of each OM system with respect to its RM counterpart, we observed a positive trend with respect to the relative increase in the volume of the parameter space in which the system was dynamically stable (Fig. 3C). Systems with consumption matrices with low connectances, although qualitatively following the same trend, departed from the mean -and RM systems showed a larger dynamically stable volume. Finally, we illustrated the increase in functional groups by representing two networks (OM and RM) (Fig. 3D). The intermediate-high connectance and consumption overlap of the network selected made apparent that, for the RM system, few functional groups having dense connections were found (represented by the proximity in space of the nodes). The optimization split these functional groups into smaller subgroups, differentiating their ecological strategies.

Our results shed some light on how complex microbial communities could form and maintain when there is syntrophy. In the comparison between systems with and without syntrophy, we chose a conservative scenario in which all resources were externally supplied. This scenario favours the feasibility of purely competitive systems, since those with syntrophy must fulfill more constraints at the fixed point, hence being less feasible. We didn’t consider other possible scenarios such as one in which some resources are not externally supplied but only produced by species, which should favour systems with syntrophy since they can exploit microniches created by microbial excretions that would not be present in the purely competitive scenario. Since, in nature, parameters are not chosen at random but they are shaped by evolution, the lower feasibility of systems with syntrophy might be irrelevant. Importantly, despite of being less feasible, syntrophic systems were more dynamically stable as quantified by the mean rate of return after a perturbation.

Among systems with syntrophy, OM systems stood out as those with a more feasible and dynamically stable volume of the parameter space. For the mean rate of return to equilibrium, however, FC systems had a superior recovery rate, but the number of feasible systems dropped for high syntrophy. Therefore, OM systems seem to stand as a compromise between feasibility and recovery rates.

Notably, we found a plausible explanation for the remarkable behaviour of OM systems through the notion of functional group. The classification of species in functional groups reflect different ecological strategies, here represented by different consumed and secreted metabolites. Increasing the number of functional groups will, therefore, reduce the ecological overlap in the resources consumed between pairs of species, which reduce the effective competition they feel. Similar results were derived with a different model in [19], in which a relationship between the existence of optimal complementary combinations of functional groups (there termed metabolic classes) and the existence of non-invadable communities (termed cartels) was found. Cartels would be consistent with the hypothesized existence of an intermediate level of organization between the single-species and multispecies populations found in natural communities, termed metabolically cohesive consortia [27], proposed to explain the remarkably high microbial biodiversity observed in nature. Indeed, functional groups complementarity would be aligned with results found in LV mutualistic systems in which it was shown that reducing interspecific competition enhanced structural stability, hence promoting biodiversity [22].

In summary, our results emphasize the importance of searching for community-level principles to understand microbial organization and its role in biodiversity. The optimization principle we found may pave the way to shed light on this question and to suggest new avenues for the optimization and design of synthetic microbial communities.

## Acknowledgements

We thank Ugo Bastolla, Michael Manhart and Justus Fink for useful discussions.

## Software availability

Software to reproduce the results presented in this work is provided in the URL: https://github.com/apascualgarcia/consumersresources

