## Supplementary Materials for "A network optimisation condition uncovers the role of functional groups in the feasibility and dynamical stability of microbial model ecosystems"

### 1 Supplementary Methods

#### 1.1 Parameterization of the model

We parameterized the model considering that each parameter $X$ followed a uniform distribution $\text{Unif}(X_0 - \delta, X_0 + \delta)$ with $\delta = 0.05X_0$. Hereafter and in Main Text, we call the parameters $X_0$ and $\delta$ metaparameters. Matrices were generated similarly, we considered $\gamma_{i\mu} = g_{i\mu} \odot G_{i\mu}$ and $\alpha_{\mu i} = a_{i\mu} \odot A_{i\mu}$, where $g_{i\mu} = \text{Unif}(\gamma_0 - \delta, \gamma_0 + \delta)$, $a_{i\mu} = \text{Unif}(\alpha_0 - \delta, \alpha_0 + \delta)$ and $G_{i\mu}(A_{i\mu})$ is a binary matrix describing the consumption (secretion) of resource $\mu$ by species $i$, and $\odot$ denotes the Hadamard product. The specific metaparameter values are summarized in Suppl. Table 1.

In our analysis, we varied three metaparameters controlling the consumption rate ($\gamma_0$), secretion rate ($\alpha_0$) and abundances at equilibrium ($S_0$). The remaining parameters were fixed as follows. The external input of resources $l_0 = 1$ and equilibrium abundances of resources $R_0 = 1$ set the units of time and biomass, respectively. To choose the efficiency of transformation of nutrients into cell biomass, $\sigma_0$, we noted (see Eq. 4 below) that the feasibility of the system depends on $\min(\sigma_0, 1 - \sigma_0)$, suggesting the existence of two regimes $\sigma_0 < 0.5$ and $\sigma_0 > 0.5$. Estimating the efficiency is a difficult task since it may depend on the environmental conditions, but it is likely small and independent of body size [1] and, therefore, we chose the interval $\sigma_0 < 0.5$ by fixing it to a constant value $\sigma_0 = 0.25$. The remaining parameters, $m_\mu$ and $d_i$ were fixed choosing a value of the variable metaparameters $\gamma_0$, $\alpha_0$ and $S_0$ and solving the system of equations at steady state.

| | Metaparameter |
|---|---|
| $S_i^*$ | $S_0 \in (0, 1]$ |
| $\gamma_{i\mu}$ | $\gamma_0 \in (0, 1]$ |
| $\alpha_{i\mu}$ | $\alpha_0 \in [0, 1]$ |
| $l_\mu$ | $l_0 = 1$ |
| $R_\mu^*$ | $R_0 = 1$ |
| $\sigma_i$ | $\sigma_0 = 0.25$ |
| $m_\mu$ | Solved at fixed point |
| $d_i$ | Solved at fixed point |

*Table 1: **Metaparameters** considered in the model.*

#### 1.2 Generation of consumption and syntrophy matrices

We explored a large number of systems with different consumption and syntrophy matrices to understand the generality of our results. Consumption matrices were generated by randomly drawing binary matrices with given connectance, at 8 uniformly-spaced values within the interval (0.05, 0.45), (red circles in Suppl. Fig. 1). Next, for each value of connectance, we modified the consumption overlap between species by applying the algorithm by Medan et al. [2] that swaps links maintaining the matrix degrees. Each swapping was selected with a Metropolis criterion that enforced convergence towards the target overlap (black circles in Suppl. Fig. 1). For each connectance value, we chose different consumption overlap values between the starting connectance (which is a lower bound for the overlap) and 0.6.

The optimized syntrophy matrices (OM) were generated as follows. For each consumption matrix, a random matrix with randomly selected connectance was drawn. Next, we applied a Monte Carlo algorithm in which, at each step, we created or destroyed a link with probability $p = 0.5$, or swapped two links following the algorithm indicated above with probability $1 - p$. When creation/destruction of a link was chosen, the two possible events happened with the same probability (0.5). We followed a Metropolis criterion to optimize the objective function $E$ until convergence, which typically happened after $4 \times 10^4$ Monte Carlo steps. However, we noted that the minimization could lead to different solutions for different runs. Therefore, to avoid biasing our results by selecting an arbitrary configuration, we run the optimization 100 times and generated a meta-matrix quantifying the frequency $p_{\mu j}$ in which a secretion of resource $\mu$ by species $j$ was observed across runs. We used this metamatrix to generate a new set of 100 syntrophy matrices by randomly drawing a random number $r$ in the interval [0, 1] for each cell, and creating a syntrophy link if $r > p_{\mu j}$. This set was the one we presented as 'optimized matrices'. Note, however, that our approach was very conservative, because the procedure likely destroyed correlations between cells that could be relevant for the properties we studied.

We further evaluated results obtained with OM systems by comparison against systems having the same connectance matrix and a fully connected syntrophy matrix (FC systems) and another set having randomly generated matrices with the same connectance than the OM found (RM systems).

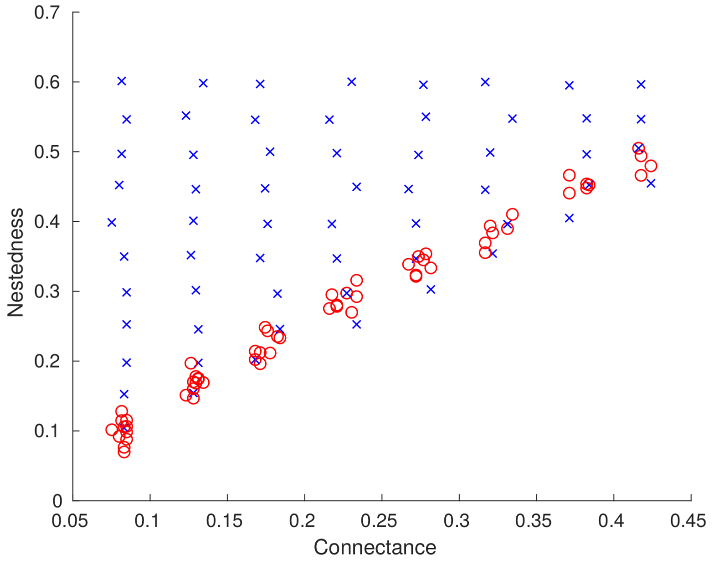

*Figure 1:* ***Generation of consumption matrices.*** *Matrices were randomly generated for different connectance values (red circles) and then randomly rearranged to attain different consumption overlap values (black circles), representing the final consumption matrices used in this study.*

#### 2 Supplementary Figures

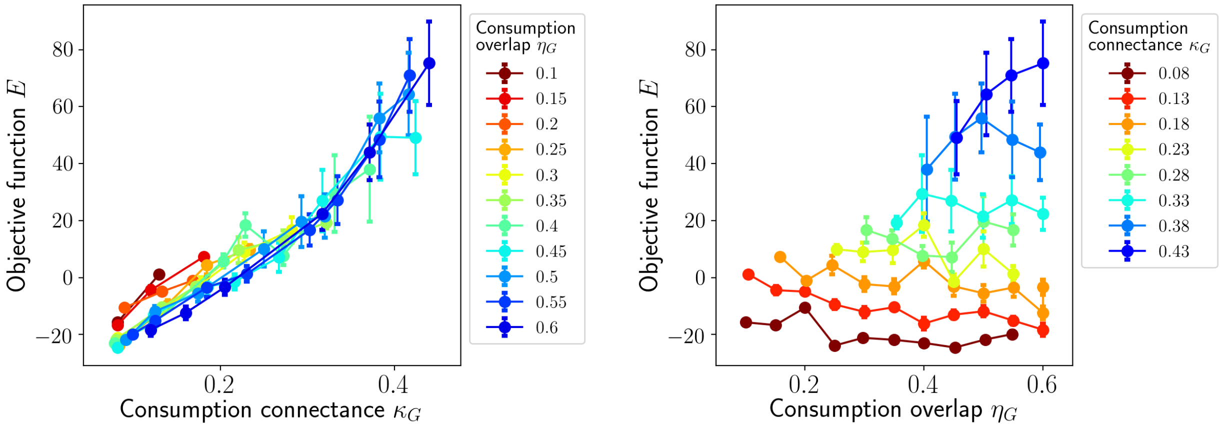

*Figure 2:* ***Objective function vs. topological properties of the consumption matrix.*** *(Left) connectance vs. objective function for fixed consumption overlap; (Right) consumption overlap vs. objective function for fixed connectance. The objective function has a strong dependence on the connectance, whereas the consumption overlap has a modest or no influence.*

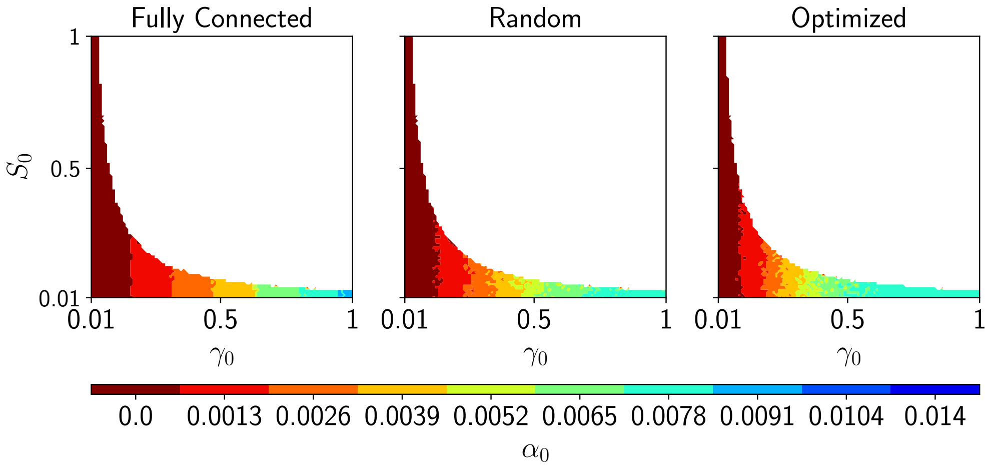

*Figure 3:* ***Computation of the feasible and dynamically stable volume.*** *Representation of the region of parameters in which a system with consumption connectance* $\kappa_G = 0.23$ *and ecological overlap* $\eta_G = 0.55$ *is feasible and dynamically stable. The same network is evaluated with different syntrophy matrices: FM (left), RM (middle) and OM (right) and for different values of syntrophy. Smaller surfaces are represented on top of larger ones, showing that larger syntrophy values make the dynamically stable volume shrink. Feasibility was verified first and, if the system was feasible, then linear dynamical stability was evaluated. Most feasible systems were dynamically stable.*

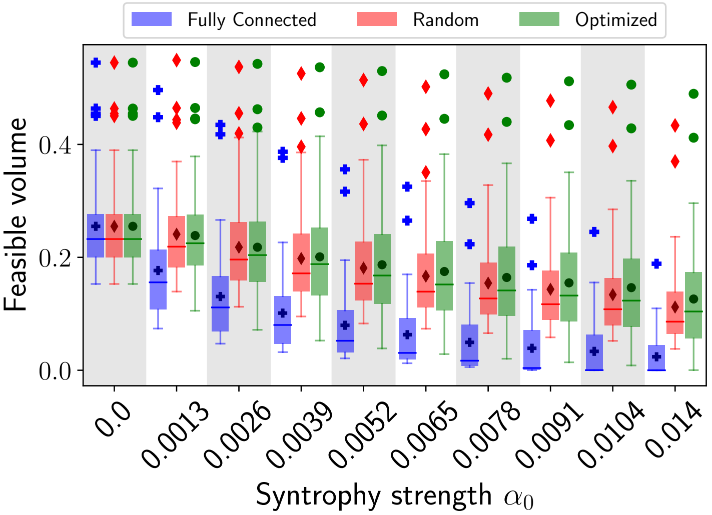

*Figure 4:* ***Feasibility volume decay.*** *Each box represents the distribution of the feasibility volume among systems with different consumption matrices and one of the three different types of syntrophy matrices. Increasing the syntrophy strength reduces the volume. The limits of each box determine the interquartile region, and the line indicates the median. Means (outliers) are represented with black(coloured) symbols.*

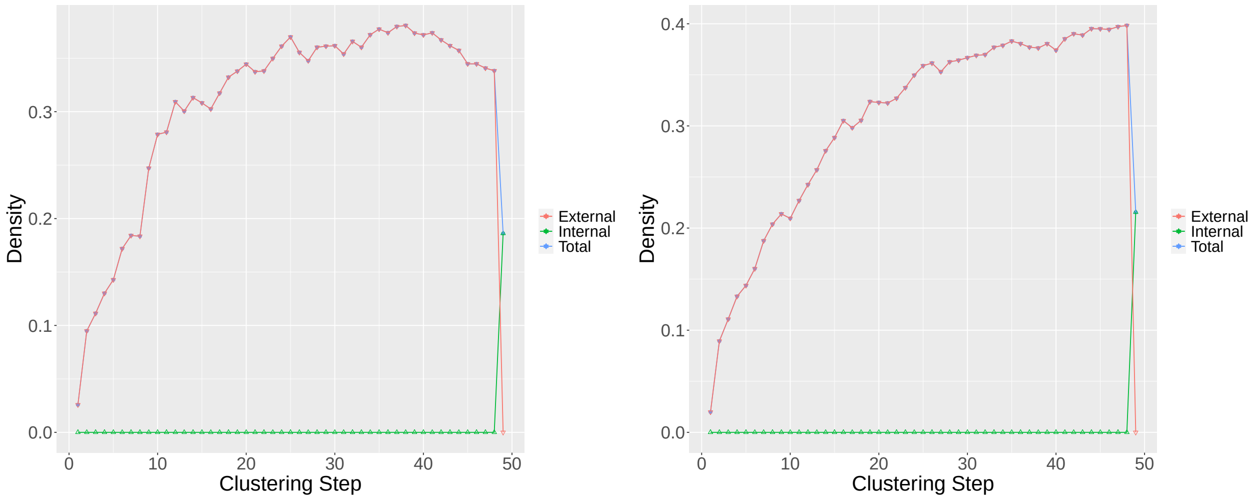

*Figure 5:* ***Determination of functional groups.*** *We searched for functional groups for each network independently. Functional groups were determined as the partition corresponding to the maximum of the total partition density (see [3] for details). Partition densities for a system with consumption connectance $\kappa_G = 0.23$ and ecological overlap $\eta_G = 0.55$ for OM (left) and RM (right) syntrophy matrices.*

#### 3 Supplementary Notes

##### 3.1 Determination of the feasibility volume

To determine the feasibility conditions we start considering the dynamical equations at a fixed point given by positive equilibrium abundances $S^*$, $R^*$:

$$
\begin{aligned}
0 &= l_\mu - \sum_j \gamma_{j\mu} S_j^* R_\mu^* + \sum_j \alpha_{\mu j} S_j^* - m_\mu R_\mu^* \\
0 &= \sum_\nu \sigma_\nu \gamma_{i\nu} R_\nu^* S_i^* - \sum_\nu \alpha_{\nu i} S_i^* - d_i S_i^*,
\end{aligned}
\tag{1}
$$

and we impose two conditions. Firstly, conservation of biomass

$$
\sum_\nu (1-\sigma_{i\nu})\gamma_{i\nu} R_\nu^* \geq \sum_\nu \alpha_{\nu i} \quad \forall i = 1, \ldots, N_{\mathrm{S}} \tag{2}
$$

and, secondly, positivity of parameters

$$
\begin{aligned}
d_i &= \sum_\nu \left(\sigma_{i\nu}\gamma_{i\nu} R_\nu^* - \alpha_{\nu i}\right) > 0 \quad \forall i = 1, \ldots, N_{\mathrm{S}} \\
m_\mu &= \frac{l_\mu - \sum_j \left(\gamma_{j\mu} R_\mu^* - \alpha_{\mu j}\right) S_j^*}{R_\mu^*} > 0 \quad \forall \mu = 1, \ldots, N_{\mathrm{R}}.
\end{aligned}
\tag{3}
$$

In the following derivation, the mean values of variables at steady state and the parameters become fixed by their respective metaparameters (see Table 1). By doing so, the most restrictive conditions imposed by Eqs. 2 and 3, will be given by the species (resources) with the most extreme matrix degrees (largest or smallest, depending on the degrees' location in the equations).

Combining conservation of biomass and positivity of parameters we find, after some algebra, that feasibility is determined by the inequalities

$$
\max_i \left\{ \frac{\deg(A,i)}{\deg(G,i)} \right\} \alpha_0 \lessapprox \min(1-\sigma_0, \sigma_0)\gamma_0 R_0 \lessapprox \min\left(1-\sigma_0, \sigma_0\right) \min_\nu \left\{ \frac{l_0}{\deg(G,\nu) S_0} + \frac{\deg(A,\nu)}{\deg(G,\nu)} \alpha_0 \right\}, \tag{4}
$$

where $\deg(X,i)$ ($\deg(X,\nu)$) indicates the degree of matrix $X$ for species $i$ (resource $\nu$) and whose limiting cases determine the more strict conditions.

In the absence of syntrophy, the second inequality in Eq. 4 becomes

$$
\gamma_0 \lessapprox \frac{l_0}{\max_\nu\{\deg(G,\nu)\} R_0 S_0}, \tag{5}
$$

indicating that the upper bound of the consumption strength is relaxed if the external rate of supplied resources increases (higher $l_0$), while it becomes more strict for larger steady state abundances ($R_0$, $S_0$) or if species consume more resources (in particular the one consuming the most, i.e. $\max_\nu\{\deg(G,\nu)\}$).

Figure 6 shows the proportion of feasible systems for 200 realizations of the parameters, for two consumption matrices $G_1$ and $G_2$. We observed a sharp transition between a region in which all realizations were feasible and another region in which all were unfeasible, with a thin transition region in which feasibility depended on the specific realization. The boundary between both regions was determined by imposing the equality in Eq. 5, which yielded $\gamma_0 = K S_0^{-1}$ with $K = l_0 / \max_\nu \{\deg(G,\nu)\} R_0$. Considering the metaparameters in the examples, we obtained $S_0 = 0.125\gamma_0^{-1}$ for $G_1$ and $S_0 = 0.077\gamma_0^{-1}$ for $G_2$, showing good agreement with a fit to the empirical values ($S_0 = (0.124 \pm 3 \times 10^{-8})\gamma_0^{-1}$ for $G_1$ and $S_0 = (0.076 \pm 7 \times 10^{-9})\gamma_0^{-1}$ for $G_2$).

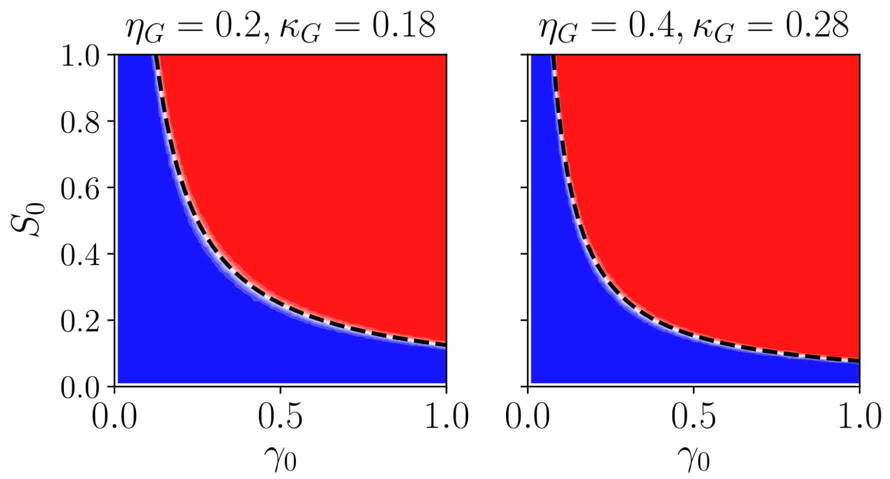

*Figure 6:* ***Analytical estimation of the feasibility boundaries.*** *Representation of the region of parameters in which two systems with different consumption connectances were feasible. The feasible (unfeasible) region is indicated in blue (red). The boundary between both regions can be derived analytically from the feasibility condition (Eq. 4), and it is represented with a dotted line.*

In Main Text, we showed that the feasibility volume shrinked when the syntrophy strength $\alpha_0$ increased. This was explained by the first inequality of Eq. 4, which shows that $\gamma_0$ is lower bounded by

$$\gamma_0 \gtrapprox \frac{\max_i \left\{ \frac{\deg(A,i)}{\deg(G,i)} \right\}}{\min(1-\sigma_0, \sigma_0) R_0} \alpha_0 .$$

As we showed when there is no syntrophy, also the abundance of consumers $S_0$ was upper bounded when there is syntrophy. Following the second inequality of Eq. 4 we found

$$\gamma_0 \lessapprox \frac{1}{R_0} \min_\nu \left\{ \frac{l_0}{\deg(G,\nu) S_0} + \frac{\deg(A,\nu)}{\deg(G,\nu)} \alpha_0 \right\},$$

which, if the resource more consumed is also the one less secreted (which would be expected e.g. if there is no intraspecific syntrophy) the analytical expression found for the case with no syntrophy would be recovered.

#### 3.2 Sufficient conditions for dynamical stability

##### The characteristic equation

To study the dynamical stability of the system we started computing its Jacobian matrix:

$$J = \begin{pmatrix} \left(-m_\mu - \sum_j \gamma_{j\mu} S_j\right) \delta_{\mu\nu} & -\gamma_{j\mu} R_\mu + \alpha_{\mu j} \\ \sigma_{i\nu} \gamma_{i\nu} S_i & \left(\sum_\nu \sigma_{i\nu} \gamma_{i\nu} R_\nu - d_i - \sum_\nu \alpha_{\nu i}\right) \delta_{ij} \end{pmatrix}, \tag{6}$$

where $\delta$ is the Kronecker delta. Since we are interested in the stability of the system at an equilibrium point $\{R^*_\mu, S^*_i\}$, from Eq. 1 we can see that

$$\sum_\nu \sigma_{i\nu}\gamma_{i\nu}R^*_\nu - d_i - \sum_\nu \alpha_{\nu i} = 0. \tag{7}$$

Hence, at equilibrium the Jacobian $J^*$ will have the following block form:

$$J^* = \begin{pmatrix} -\mathrm{D} & \Gamma \\ \mathrm{B} & 0 \end{pmatrix}, \tag{8}$$

where

- $D_{\mu\nu} = \mathrm{diag}(m_\mu + \sum_j \gamma_{j\mu}S^*_j) = \mathrm{diag}\left(\frac{l_\mu + \sum_j \alpha_{\mu j}S^*_j}{R^*_\mu}\right)$ is a positive $N_R \times N_R$ diagonal matrix,
- $\Gamma_{\mu j} = -\gamma_{j\mu}R^*_\mu + \alpha_{\mu j}$ is a $N_R \times N_S$ is a matrix which does not have entries with a definite sign.
- $B_{i\nu} = \sigma_{i\nu}\gamma_{i\nu}S^*_i$ is a $N_S \times N_R$ is a matrix with positive entries.

The stability of the system was analysed studying the eigenvalue problem determined by the characteristic equation

$$\det\left(J^* - \lambda\right) = 0 \tag{9}$$

or, more explicitly,

$$\det\begin{pmatrix} -\mathrm{D} - \lambda & \Gamma \\ \mathrm{B} & -\lambda \end{pmatrix} = 0. \tag{10}$$

If $\lambda \neq 0$, it is possible to simplify this equation using the properties of block matrices [4]:

$$\det\begin{pmatrix} -\mathrm{D} - \lambda \mathrm{I}_{N_R} & \Gamma \\ \mathrm{B} & 0 - \lambda \mathrm{I}_{N_S} \end{pmatrix} = \det\left(-\lambda \mathrm{I}_{N_S}\right)\det\left(-\mathrm{D} - \lambda \mathrm{I}_{N_R} + \frac{1}{\lambda}\Gamma\mathrm{B}\right). \tag{11}$$

Hence, Eq. (10) becomes:

$$\det\left(\lambda^2 \mathrm{I}_{N_R} + \mathrm{D}\lambda - \Gamma\mathrm{B}\right) = 0. \tag{12}$$

Note that the complexity was reduced because we went from the determinant of a $N_R + N_S$ square matrix to a $N_R$ square matrix. Using standard properties of determinants, we can rewrite Eq. (12) as[1]:

$$\det\left(-\mathrm{D}\right)\det\left(-\mathrm{D}^{-1}\lambda^2 - \lambda + \mathrm{D}^{-1}\Gamma\mathrm{B}\right) = 0 \iff \det\left(S(\lambda) - \lambda\right) = 0 \tag{13}$$

with

$$S(\lambda) = \mathrm{D}^{-1}\Gamma\mathrm{B} - \mathrm{D}^{-1}\lambda^2, \tag{14}$$

or, component-wise,

$$S_{\mu\nu} = \frac{1}{D_\mu}\left[\left(\sum_i \mathit{\Gamma}_{\mu i}B_{i\nu}\right) - \lambda^2\delta_{\mu\nu}\right], \tag{15}$$

where the $\Gamma\mathrm{B}$ $N_R$-dimensional square matrix is given by:

$$(\mathit{\Gamma}B)_{\mu\nu} = \sum_i \mathit{\Gamma}_{\mu i}B_{i\nu} = \sum_i \left(\alpha_{\mu i} - \gamma_{i\mu}R^*_\mu\right)\sigma_{i\nu}\gamma_{i\nu}S^*_i. \tag{16}$$

The strategy we used to solve Eq. (13) was inspired by the one followed in Ref. [5]. We assumed that, if the system is in an unstable regime, *i.e.* there exists at least one $\lambda \in \sigma(J^*)$ satisfying Eq. (12) such that $\mathrm{Re}\left(\lambda\right) > 0$. By Eq. (13), $\lambda$ is also an eigenvalue of $\mathrm{S}(\lambda)$. We reasoned that, if we find conditions in which the real part of the spectrum of $\mathrm{S}(\lambda)$ is entirely negative, we will know that $\mathrm{Re}\left(\lambda\right) \leq 0$. As this is a contradiction to the hypothesis that the regime is unstable, we must conclude that the regime is stable[2]. Hence, the general idea was to find regimes where we knew that the spectrum of $S$ was entirely negative for a positive $\lambda$.

[1]Since $m_\mu > 0$, we know $D$ will always be invertible.

[2]Indeed, Eq.(12) assumes already that either $\mathrm{Re}\left(\lambda_1\right) > 0$ or $\mathrm{Re}\left(\lambda_1\right) < 0$.

**Derived conditions from Gerschgorin circle theorem**

The Gerschgorin circle theorem [6] states that every eigenvalue of a $N \times N$ square matrix A is located in one of the $N$ discs $\tilde{D}_i$ defined by:

$$\tilde{D}_i \equiv \left\{ z \in \mathbb{C} : |z - A_{ii}| \leq \sum_{j \neq i} |A_{ij}| \right\}. \tag{17}$$

Intuitively, the circle theorem tells us that the eigenvalues of a matrix deviate from the diagonal elements by a value bounded by the sum of the off-diagonal elements. We noted that, if all the discs $\tilde{D}_i$ are located to the left of the imaginary axis (*i.e.* the discs contain only numbers with a negative real part), then the eigenvalues of A are all negative. This theorem allowed us to consider two results in our problem. Firstly, it provided the following bound for the eigenvalues

$$|\lambda| \leq r_{\mathrm{c}} \quad \forall \lambda \in \sigma(J^*), \tag{18}$$

where $r_{\mathrm{c}}$ is the*critical radius* and $\sigma(J^*)$ stands for the spectrum of the Jacobian at equilibrium. To estimate $r_{\mathrm{c}}$ we noted that all eigenvalues of $J^*$ were located in one of the $N_{\mathrm{R}} + N_{\mathrm{S}}$ discs of $J^*$, among the $N_{\mathrm{R}}$ "resources discs":

$$\tilde{D}^{\mathrm{R}}_{\mu} \equiv \left\{ z \in \mathbb{C} : |z + D_\mu| \leq \sum_{j} |\Gamma_{\mu j}| \right\} \ \forall \mu = 1, \ldots, N_{\mathrm{R}}, \tag{19}$$

and the "consumers discs":

$$\tilde{D}^{\mathrm{C}}_{i} \equiv \left\{ z \in \mathbb{C} : |z| \leq \sum_{\nu} |B_{i\nu}| \right\} \ \forall i = 1, \ldots, N_{\mathrm{S}}. \tag{20}$$

According to the circle theorem Eq.(17), all eigenvalues are in the union of these circles, *i.e.* there exists $\forall \lambda \in \sigma(J^*)$ at least one $\mu^*$ or one $i^*$ such that:

$$|\lambda| \leq \sum_{\nu} |B_{i^* \nu}| \tag{21}$$

or

$$|\lambda + D_{\mu^*}| \leq \sum_{j} |\Gamma_{\mu^* j}|. \tag{22}$$

The triangle inequality implies:

$$|\lambda| \leq |\lambda + D_{\mu^*}| + |-D_{\mu^*}| \leq \sum_{j} |\Gamma_{\mu^* j}| + |-D_{\mu^*}| = \sum_{j} |\Gamma_{\mu^* j}| + D_{\mu^*}. \tag{23}$$

The only way both Eq.(21) and (23) are satisfied for all eigenvalues, and especially the one with the highest real part $\lambda_1$ is if they are bound by the maximum of both RHS of these equations, which allowed us to determine $r_{\mathrm{c}}$ as:

$$r_{\mathrm{c}} \equiv \max \left\{ \max_{i} \left\{ \sum_{\nu} |B_{i\nu}| \right\}, \max_{\mu} \left\{ \sum_{j} |\Gamma_{\mu j}| + D_\mu \right\} \right\}. \tag{24}$$

The second result derived from the theorem is the following Lemma, which puts an upper bound on the real part of the spectrum[3] of any square matrix.

**Lemma 1.** *If a $N-$dimensional square matrix $A$ verifies the equations:*

$$Re(A_{ii}) + \sum_{j \neq i} |A_{ij}| < 0, \forall \quad i = 1, \ldots, N, \tag{25}$$

*then* $Re(\lambda) < 0 \quad \forall \lambda \in \sigma(A)$.

[3]We denote the spectrum of a matrix $M$ with the symbol $\sigma(M)$.

*Proof.* Let $\lambda \in \sigma(A)$. By the circle theorem, there exists $k \in \{1, \dots, N\}$ such that :

$$|\lambda - A_{kk}| \leq \sum_{j \neq k} |A_{kj}| \,. \tag{26}$$

We now use the complex identity:

$$|\lambda - A_{kk}| \geq \mathrm{Re}\,(\lambda - A_{kk}) = \mathrm{Re}\,(\lambda) - \mathrm{Re}\,(A_{kk}) \,. \tag{27}$$

Equation (25) implies:

$$\sum_{j \neq k} |A_{kj}| < -\mathrm{Re}\,(A_{kk}) \,. \tag{28}$$

Combining the two previous inequalities yields:

$$\mathrm{Re}\,(\lambda) - \mathrm{Re}\,(A_{kk}) \leq |\lambda - A_{kk}| \leq \sum_{j \neq k} |A_{kj}| < -\mathrm{Re}\,(A_{kk}) \,. \tag{29}$$

Comparing the RHS and LHS of this inequality yields:

$$\mathrm{Re}\,(\lambda) < 0. \tag{30}$$

□

**Sufficient condition for dynamical stability (strong)**

**Theorem 1.** *Let $p$ be a parameter set with a Jacobian at equilibrium $J^*$. If $0$ is not an eigenvalue of $J^*$ and the equations*

$$(\mathit{\Gamma} B)_{\mu\mu} < -\sum_{\nu \neq \mu} \left|(\mathit{\Gamma} B)_{\mu\nu}\right| - r_{\mathrm{c}}^2 \ \forall \mu, \tag{31}$$

*are verified, then $p$ is dynamically stable.*

*Proof.* We assume

$$(\mathit{\Gamma} B)_{\mu\mu} < -\sum_{\nu \neq \mu} \left|(\mathit{\Gamma} B)_{\mu\nu}\right| - r_{\mathrm{c}}^2 \ \forall \mu. \tag{32}$$

This implies:

$$(\mathit{\Gamma} B)_{\mu\mu} + r_{\mathrm{c}}^2 < -\sum_{\nu \neq \mu} \left|(\mathit{\Gamma} B)_{\mu\nu}\right| \ \forall \mu. \tag{33}$$

If the previous inequality is verified, since $\mathrm{Re}\,(\lambda)^2 \leq |\lambda|^2 < r_{\mathrm{c}}^2$, the following one is also trivially verified,

$$(\mathit{\Gamma} B)_{\mu\mu} - \mathrm{Re}\,(\lambda)^2 < -\sum_{\nu \neq \mu} \left|(\mathit{\Gamma} B)_{\mu\nu}\right| \ \forall \mu. \tag{34}$$

Using this result and dividing Eq.(34) by $D_\mu$ ($D_\mu > 0$, $\forall \mu$), we get:

$$\frac{1}{D_\mu}\left[\left(\sum_i \mathit{\Gamma}_{\mu i} B_{i\mu}\right) - \mathrm{Re}\,(\lambda^2)\right] < -\sum_{\nu \neq \mu} \left|\frac{\sum_i \mathit{\Gamma}_{\mu i} B_{i\nu}}{D_\mu}\right| \ \forall \mu. \tag{35}$$

Looking at Eq.(15), we see that Eq.(35) is equivalent to:

$$\mathrm{Re}\,(S_{\mu\mu}) + \sum_{\nu \neq \mu} |S_{\mu\nu}| < 0 \ \forall \mu. \tag{36}$$

Lemma 1 implies that all the eigenvalues of $S(\lambda)$ have a negative real part. By *reductio ad absurdum,* if $\mathrm{Re}\,(\lambda) \geq 0$ in Eq. (14) (unstable or marginally stable regime), then $\mathrm{Re}\,(\lambda) < 0$ in Eq. (35), which leads to a contradiction. This then implies that the equilibrium is dynamically stable. □

Note that Theorem 1 is a sufficient condition, and there may be dynamically stable systems that do not fulfill it. It demands strong constraints on the parameters set, namely ΓB must have negative diagonal elements, imposing constraints on the $\alpha$ and $\gamma$ matrices.

##### Sufficient condition for dynamical stability (weak)

Another theorem can be stated with less restrictive assumptions which, however, leads to a less general condition.

**Theorem 2.** *Let $p$ be a parameter set with a Jacobian at equilibrium $J^*$. If 0 is not an eigenvalue of $J^*$ and the equations*

$$(\mathit{\Gamma} B)_{\mu\mu} < -\sum_{\nu\neq\mu} \left|(\mathit{\Gamma} B)_{\mu\nu}\right| \;\forall\mu, \tag{37}$$

*are verified, then the real eigenvalues of $J^*$ are negative.*

*Proof.* We assume

$$(\mathit{\Gamma} B)_{\mu\mu} < -\sum_{\nu\neq\mu} \left|(\mathit{\Gamma} B)_{\mu\nu}\right| \;\forall\mu. \tag{38}$$

Let $\lambda \in \mathbb{R}$, then the following will also trivially hold:

$$(\mathit{\Gamma} B)_{\mu\mu} - \lambda^2 < -\sum_{\nu\neq\mu} \left|(\mathit{\Gamma} B)_{\mu\nu}\right| \;\forall\mu. \tag{39}$$

Dividing Eq.(39) by $D_\mu$, we get:

$$\frac{1}{D_\mu}\left[\left(\sum_i \mathit{\Gamma}_{\mu i} B_{i\mu}\right) - \lambda^2\right] < -\sum_{\nu\neq\mu} \left|\frac{\sum_i \mathit{\Gamma}_{\mu i} B_{i\nu}}{D_\mu}\right| \;\forall\mu. \tag{40}$$

Looking at Eq.(15), we see that this is equivalent to:

$$S_{\mu\mu} + \sum_{\nu\neq\mu} |S_{\mu\nu}| < 0 \;\forall\mu. \tag{41}$$

Using Lemma 1, we know that all the real eigenvalues of §($\lambda$) will have a negative real part. We can conclude with the statement of the theorem. □

The condition is less general because it only applies for real eigenvalues of $J^*$, hence excluding complex eigenvalues.

##### Objective function derivation

The two theorems derived suggested that dynamical stability would more likely be fulfilled if the matrix ΓB is diagonally dominant, i.e. if the condition in Eq. (38) is fulfilled. Therefore, we proposed an objective function whose minimization allowed us to search for topologies more likely leading to dynamically stable systems

$$E = \sum_\mu \left((\mathit{\Gamma} B)_{\mu\mu} + \sum_{\nu\neq\mu} \left|(\mathit{\Gamma} B)_{\mu\nu}\right|\right)$$

which we can make more explicit in terms of the metaparameters and binary matrices, to get the expression presented in Main Text:

$$E(A,G) = \sum_\mu (\alpha_0 \mathrm{AG} - \gamma_0 R_0 \mathrm{G}^T\mathrm{G})_{\mu\mu} + \sum_{\mu\neq\nu} \left|(\alpha_0 \mathrm{AG} - \gamma_0 R_0 \mathrm{G}^T\mathrm{G})_{\mu\nu}\right|.$$
